# Identification and functional analysis of novel stress-resistance genes from metagenomes of extreme environments

**DOI:** 10.1101/2023.06.07.544099

**Authors:** Joshelin Huanca Juarez, Edson do Nascimento Silva, Ninna Hirata Silva, Rafael Silva-Rocha, María-Eugenia Guazzaroni

## Abstract

Currently, industrial bioproducts are less competitive than chemically produced goods due to the shortcomings of conventional microbial hosts. Metagenomic approaches from extreme environments can provide useful biological parts to improve bacterial robustness to process-specific parameters. Here, in order to build synthetic genetic circuits that increase bacterial resistance to diverse stress conditions, we mined novel stress tolerance genes from metagenomic databases using an *in silico* approach based on Hidden-Markov-Model profiles. For this purpose, we used metagenomic shotgun sequencing data from microbial communities of extreme environments to identify genes encoding chaperones and other proteins that confer resistance to stress conditions. We identified and characterized ten novel protein-encoding sequences related to the DNA-binding protein HU, the ATP-dependent protease ClpP, and the chaperone protein DnaJ. By expressing these genes in *Escherichia coli* under several stress conditions (including high temperature, acidity, oxidative and osmotic stress, and UV radiation), we identified five genes conferring resistance to at least two stress conditions when expressed in *E. coli*. Moreover, one of the identified HU coding-genes which was retrieved from an acidic soil metagenome increased *E. coli* tolerance to four different stress conditions, implying its suitability for the construction of a synthetic circuit directed to expand broad bacterial resistance.

## Introduction

Industrial biotechnology focuses on working with biological systems, such as cell factories or enzymes, rather than using fossil raw materials to produce antibiotics, chemicals, and energy sources (Festel, 2020). As a result, it has attracted much attention for its environmentally friendly and sustainable methods, which could pave the way for a paradigm shift from petroleum to bio-based production (L.-P. Yu et al., 2019). Therefore, this industry is positioned as a solution to critical global concerns such as climate change mitigation, food security, ocean pollution reduction, and biodiversity conservation (Rosemann & Molyneux-Hodgson, 2020). Microbial Cell Factories, however, are subjected to harsh extracellular conditions such as high temperature, low pH, and metabolite toxicity, resulting in lower productivity than generated on a laboratory scale and, consequently, higher production costs (Gong et al., 2017; Jiang et al., 2020). Highly robust and tolerable strains towards non-natural or even toxic substrates or products and abiotic conditions are required since they can utilize cheaper carbon and energy sources, such as waste residues or biomass hydrolysates (Blombach et al., 2022).

Abiotic stress conditions – such as those present in industrial bioprocesses – produce considerable cellular damage in non-adapted microorganisms, affecting the structure and stability of various biomolecules, including nucleic acids, membrane lipids, and proteins (Merino et al., 2019; Rothschild & Mancinelli, 2001). To counteract these damage conditions, cells activate different repair mechanisms, such as DNA/RNA repair proteins, proteases to remove unfolded proteins, and multifunctional enzymes, such as RecA, Hup, RecR, RecO, PriA, and RecN. These enzymes are involved in DNA condensation, remotion of broken DNA, and DNA recombination repair (Grosjean & Oshima, 2007; Somayaji et al., 2022; G. Wang et al., 2011). There is also a subset of activated post-translationally protein under specific stress conditions known as molecular chaperones, which act in several processes involving cellular protein folding, unfolding, and homeostasis (Saibil, 2013; Voth & Jakob, 2017). Considering that protein denaturation is a persistent direct or indirect result of stress, chaperones are vital in defensive cellular mechanisms (Jacob et al., 2017).

In addition to the group of enzymes mentioned above, there are other proteins of particular relevance as protective biological agents against stress damage to essential macromolecules. For instance, the DNA-binding protein HU (histone-like protein) has been shown to bind with high specificity to aberrant DNA formations such as nicks and strand breaks to preserve its integrity (Kamashev & Rouviere-Yaniv, 2000). The Clp protease complex consists of a proteolytic component, ClpP, in association with any of the ATPases, ClpA or ClpX, granting substrate specificity (Khan et al., 2022; J. Wang et al., 1997). Proteases, in general, play an essential role in protein quality control by removing short-lived regulatory proteins as well as misfolded and damaged proteins, which are toxic to the cell (A. Y. H. Yu & Houry, 2007). Lastly, DnaJ, also known as J-protein, is a homolog of eukaryotic HSP40 and acts as a cofactor of DnaK, stimulating its ATPase activity through the J domain, while additional domains give specificity to the system (Cheetham & Caplan, 1998; Ghafoori et al., 2017; Kampinga et al., 2019). In addition to participating in other metabolic tasks, these molecular chaperones have been shown to play an essential role in protein degradation (Huang et al., 2001).

In recent years, various studies have focused on the capacity of bacteria to survive in extreme environments in order to understand the mechanisms of biochemical adaptation to harsh conditions (Guazzaroni et al., 2015; Mirete et al., 2016; Morozkina et al., 2010). Extremophiles live in a wide range of hostile conditions and have evolved a set of adaptations to cope with these situations (Coker, 2019). However, most of them are not cultivable, which limits their exploration in biotechnological applications (Alves, Westmann, et al., 2018; Rappé & Giovannoni, 2003). Culture-independent approaches help harness the biological activity and build components of this difficult-to-grow biodiversity (van der Helm et al., 2018). In this context, functional and sequence-based metagenomics have been shown to be effective in the identification of novel genes that confer resistance to extreme conditions, such as enzymes, antibiotics, and other bioactive molecules derived from a variety of environments (Alves, Meleiro, et al., 2018). Functional metagenomic approaches allowed to identify novel genes relevant to biotechnology, involving the creation of expression libraries with thousands of metagenomic clones and activity-based screens in order to find genes that encode a function of interest (Guazzaroni et al., 2013; Mirete et al., 2016). On the other hand, the search for sequences of interest can also be done by sequence identification in (meta)genomic databases; however, low-level expression of heterologous proteins in commonly used hosts and low quality of (meta)genomic annotations are the main disadvantages in both approaches, respectively (Alves, Westmann, et al., 2018; van der Helm et al., 2018). Furthermore, sequence-based screenings generally employ sequence pair aligners (such as BLAST, BLAT, Minimap2, KMA, and Bowtie) to detect regions of similarity between two biological sequences (Altschul et al., 1990; Clausen et al., 2018; Kent, 2002; Langmead & Salzberg, 2012). These tools constrain the homologous detection process since the percentage of identity is superior to a certain threshold, so more distant evolutionary sequences are lost (Skewes-Cox et al., 2014).

On the other hand, computational methods based on profiles, such as Hidden Markov Model (HMM), allow the detection of remote protein homologs (Kirsip & Abroi, 2019; Radivojac, 2022). In this context, profiling methods gained sensitivity by incorporating position-specific information into the alignment process and quantifying variation between family members at each position (Madera & Gough, 2002; Skewes-Cox et al., 2014). Some software based on HMM profiles were shown to be useful for identifying sequences of interest, such as IDOP or CryProcessor, used for detecting toxin genes (Díaz-Valerio et al., 2021; Shikov et al., 2020). In the same way, deep learning has been shown to be a powerful machine learning approach in predicting DNA sequence affinities and identifying new genes of interest, such as resistance genes or transcription factors (Arango-Argoty et al., 2018; Oliveira Monteiro et al., 2022). Considering the above, we aimed to identify microbial genes related to stress resistance by mining sequences from available metagenomic datasets from extreme environments using HMM profiles. For this achievement, we focused on searching novel protein sequences associated with the DNA-binding protein HU, the ATP-dependent protease ClpP, and the chaperone protein DnaJ. Then, we characterized their functional performance in *Escherichia coli* under five stress conditions (high temperature, acidity, oxidative and osmotic stress, and UV radiation). Through this *in silico* approach, we identified novel genes with the capacity to expand resistance to different stress conditions in bacteria, which have a high potential for the construction of a synthetic circuit directed to expand broad bacterial resistance.

## Methodology

### Bacterial strains and culture conditions

The bacterial strain *E. coli* DH10B was employed as a host in the stress resistance experiments. It was routinely grown in M9 minimal medium (1x M9 salts, 2 mM MgSO_4_, 0.1 mM CaCl_2_, 0.1% casamino acids, 1% glycerol) under aerobic conditions at 37 °C. Liquid cultures were cultivated in a shaker at 37°C and 200 rpm. For plasmid maintenance, the growth medium for bacteria transformed with the synthetic circuits was supplied with 150 μg/mL ampicillin (Ap). For stress assays, overnight cultures were diluted to a final OD_600_ 0.05 in 5 mL of M9 with antibiotic and incubated in an orbital shaker at 37°C and 200 rpm. When cultures reached the early exponential growth phase (OD_600_ = 0.3 - 0.4), stress tests were initiated.

### Selection of metagenomes from public databases

Raw shotgun metagenomes were curated, chosen, and downloaded from the MG-RAST database, hosted by the University of Chicago and the Argonne National Laboratory (https://www.mgrast.org/). The data were then pre-processed using FastQC to check the sequence’s quality, and when necessary, low-quality sequences were excluded using the Trimmomatic tool (Bolger et al., 2014). After that stage, metagenomes were assembled into contigs using the MEGAHIT program (D. Li et al., 2015), and an initial annotation was performed using Prokka.

### Selection of genes of interest and construction of HMM profiles

HMM profiles were built based on selected stress-resistance genes previously documented in the literature. For this, we selected genes of nucleic acid binding protein genes (RBP, HU, DPS); DNA replication protein genes (GyrA, RecA, DnaA), chaperone and protease genes (ClpP, ClpX, ClpA, ClpC, ClpE, ClpL, DnaJ, DnaK); and CRISPR system genes (Cas1, Cas2, Cas9). The nucleotide sequences were obtained from the National Center for Biotechnology Information (NCBI) database, and alignment was performed with MAFFT (Multiple Alignment using Fast Fourier Transform) (Katoh et al., 2002). The profiles were created using the HMMer3 tool (Eddy, 2009), which was also used to detect homologous sequences by comparing the HMM profiles generated and the sequences of previously selected metagenomes. The results were analyzed and correlated with the literature, aiming to obtain ten candidate genes. The selection process of protein sequences for further functional characterization was based on the following characteristics: (1) abundance of the proteins in the different environments, (2) metagenomic genes related to stress resistance in the literature, and (3) protein function. To continue the analysis and selection of sequences, a BLASTp was performed, considering an identity percentage threshold of less than 80% (https://blast.ncbi.nlm.nih.gov).

### *In silico* analysis and 3D-Structure Model Generation

Dendrograms for each class of protein were constructed based on the 23-25 amino acid sequences using MEGA X software. The homologous sequences of selected proteins in different organisms from the NCBI database were retrieved in FASTA format. Alignment was done using the Muscle algorithm (Edgar, 2004) and used to build the dendrogram. The Neighbor-joining statistical method was applied, and the accuracy of the phylogenetic analysis was predicted by using 1,000 bootstrapping replications. The 3D-structure models were generated using the Alphafold2 collab server (Jumper et al., 2021). All structures were analyzed using the PyMol v. 2.5.2 software (http://www.pymol.org/) where the RMSD value of the alpha carbon atoms was considered. Proteins were aligned with T-Coffee v. 13.45.60.cd84d2a, and the alignment results were visualized in Jalview v. 2.11.2.3.

### Cloning of the selected genes

All amino acid sequences previously selected by bioinformatics analysis were converted to nucleotides sequences using the EMBOSS Backtranseq tool (https://www.ebi.ac.uk/Tools/st/emboss_backtranseq/). Each nucleotide sequence was cloned under control from the constitutive promoter BBBa_J23106 (*Pj106*) and the ribosome binding site (RBS) BBa_B0034 (Mahajan, 2003). Promoter and RBS sequences were obtained from the iGEM Foundation Standard Biological Parts Catalog (Registry) (http://parts.igem.org). The genetic circuit was assembled into the pUC19 vector. The synthesis of all plasmids containing the promoter, RBS, and candidate genes cloned into the pUC19 vector was requested from the company Integrated DNA Technologies (IDT). *E. coli* DH10B competent cells were transformed with the resulting plasmids.

### Growth rate measurement and fitness cost calculation

Cell growth was monitored using a Victor X3 plate reader (PerkinElmer, Inc), as described by (de Siqueira et al., 2020). Microbial cultivation was evaluated at 30 ºC for 8 h. All experiments were performed with four biological replicates. for fitness cost calculation we followed the methodology described by (de Siqueira et al., 2020).

### Heat shock experiments

Early exponential phase cultures of *E. coli* DH10B (OD_600_ = 0.3 - 0.4) carrying an empty plasmid or the pUC19 vector with *hup, clpP*, or *dnaJ* genes were diluted at 10^−1^ in PBS buffer (pH 7.2). Then, dilutions were incubated at 55 ºC, and aliquots were removed at 0 and 30 min. To determine CFU/mL, serial dilutions were made, and three 25 μL droplets (technical replicates) were placed onto M9 plates with Ap. Plates were incubated at 37°C for 24 h. Survival percentage was expressed as the number of CFU/mL remaining after heat treatment divided by the initial CFU/mL. Each experiment was performed at least three times.

### Acid resistance assay

Early exponential phase cultures of *E. coli* DH10B (OD_600_ = 0.3 - 0.4) carrying an empty plasmid or the pUC19 vector with *hup, clpP*, or *dnaJ* genes were diluted at 10^−1^ in PBS buffer (pH 7.2) or M9 acidic medium (pH = 3.0) and then incubated at 37 ºC. Aliquots were taken at 0 and 60 min, respectively. To determine CFU/mL, serial dilutions were made as previously described. Survival percentage was the number of CFU/mL remaining after acidity treatment divided by the CFU/mL at zero time. Each experiment was repeated at least three times.

### Oxidative stress experiments

Early exponential phase cultures of *E. coli* DH10B (OD_600_ = 0.3 - 0.4) carrying an empty plasmid or the pUC19 vector with *hup, clpP*, or *dnaJ* genes were diluted at 10^−1^ in PBS buffer (pH 7.2) or M9 + hydrogen peroxide 12 mM and incubated at 37 ºC. Aliquots were taken at 0 and 60 min, respectively. Survival percent was calculated as above. Each experiment was repeated at least three times.

### UV radiation experiments

Overnight cultures of *E. coli* DH10B carrying an empty plasmid or a pUC19 vector with *hup, clpP*, or *dnaJ* genes were diluted in PBS buffer to 1.0 OD_600_. For UV radiation exposure, 10 μL of each culture were linearly spread onto M9 plates and irradiated with a germicidal lamp for 0, 5, 10, 15, 20, and 25 s at room temperature. Bacterial growth was evaluated after 24 h of incubation at 37°C. Each experiment was repeated at least three times.

### Osmotic stress assay

For continuous NaCl exposure experiments, cells were cultivated in M9 medium + 3.5% NaCl. Cultures were placed in a 96-well plate and incubated at 30°C for 8 h, with spot measurements being made every 30 minutes using a Victor X3 plate reader (PerkinElmer, Inc). All experiments were performed with four biological replicates.

### Data analysis and visualization

Data analysis and generation of growth curves were performed using the R statistical platform (version 4.2.0), using packages *ggpubr* and *ggplot2*. The *mipreadr* package version 0.1.0 was used to import and process the raw data produced by the plate reader. A Student’s t-test with a 5% significance level was employed to compare the values obtained. For clustering construction, we provide discrete arbitrary values to the different stress responses presented by clones for comparative purposes (**Table S7**). Thus, data were classified as high, medium, and low-stress responses, and values of 2, 1, and 0 were given, respectively. Then, hierarchical clustering and normalization of data were performed using the R statistical platform (version 4.2.0).

## Results and discussion

### *In silico* screening of stress-related genes

The search for new stress-resistance genes is a useful way for expanding robustness in microorganisms of industrial interest (G.-Q. Chen & Jiang, 2018; Wehrs et al., 2019). Although metagenomics offers an excellent alternative for discovering novel functional genes, activity-driven screenings are time-consuming and present low recovery rates (Guazzaroni et al., 2015; van der Helm et al., 2018). On the other hand, sequence-driven screening depends on the quality of the sequences – which present high annotation errors – and does not allow the discovery of new sequences if they have not been previously related to a specific function (Alves, Westmann, et al., 2018). In turn, profile search methods are more sensitive than pairwise alignment in detecting low percent identity homologs by incorporating position-specific information into the alignment process and quantifying variation across family members at each position (Madera & Gough, 2002; Skewes-Cox et al., 2014). Considering that, we addressed the identification of new genes in metagenomic databases using an *in silico* approach based on HMM profiles. The general computational pipeline began searching for metagenomic shotgun sequencing data from seven microbial communities of extreme environments recruited from the MG-RAST database. Then, we aimed to identify genes encoding chaperones and other proteins – such as proteases and nucleic acid-binding proteins – that confer resistance to stress conditions (**Figure 1A**).

**Figure 1.**
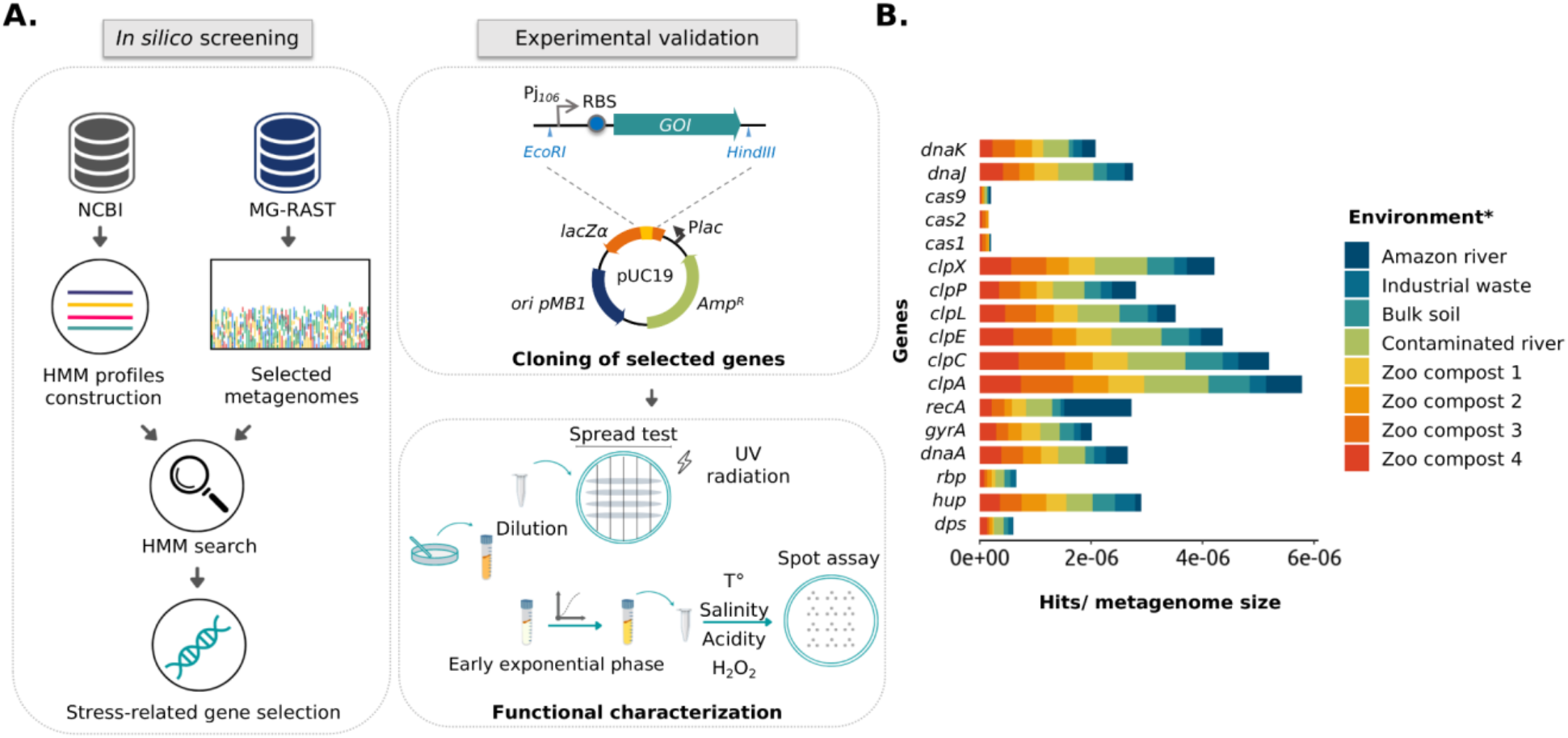
Schematic representation of the approach used in this study and frequency of stress-related proteins in public metagenomic databases. (A) Schematic representation of *in silico* and experimental approaches used to identify and validate stress-related genes using HMM profiles, metagenomic mining, and experimental characterization. (B) Representation of gene frequency in different metagenomes selected from public databases. Frequency calculation was made by normalizing the number of found hits by the size of the metagenomic library (Gb). *Metagenomes from the Atacama Desert and Dourados are not presented in the B panel, since these metagenomic data did not include all classes of the proteins plotted.

As the first step for selecting the metagenomes, we selected those who meet the following requirements: (i) that were sequenced through the Illumina platform; (ii) that the sequences did not belong to DNA from the microbiota of humans/animals; (iii) that the size of the studies was around 1 gigabyte of DNA data, and; (iv) that metagenomes were originated from environments presenting harsh conditions (or in which there was some marked abiotic stress). Selected metagenomes are listed in **Table S1**, it should be noted that about 85% are from the Brazilian territory. A summary of the metric values returned, and the annotation results are shown in **Tables S2** and **S3**, respectively. Next, to create an HMM profile, a multiple sequence alignment of 17 different stress-related proteins deposited in the NCBI database was performed using the HMMer3 server (**Figure S1**). Then, metagenome data were searched against those HMM probabilistic profiles. The resulting profiles were compared with hypothetical and non-hypothetical regions of annotated regions generating a total of 6840 hits (**Table S4**).

For the selection of protein sequences, it was considered that a protein found in a broader range of environments would be linked to the response to more than one stress factor. **Figure 1B** shows the abundance of the 17 protein families in the assembled metagenomes, considering the number of hits normalized by the size of the corresponding metagenome. The graph shows that the *rbp* and *dps* genes had the lowest abundance, being disregarded in the subsequent analysis. Then, we carried out an extensive literature search and found that 50-70% of the recovered published works connected *hup, clpP, clpX, dnaK*, and *dnaJ* genes to stress-defense mechanisms in microbial cells. As mentioned before, ClpP and ClpX protease subunits were reported to work together to produce a complex that degrades misfolded proteins and protects cells against its toxic effect (Baker & Sauer, 2012). However, different studies demonstrated that the ClpP protease subunit can function independently of the ATPase subunit ClpX (de Siqueira et al., 2020; Guazzaroni et al., 2013). On the other hand, the overexpression of the DnaK protein in *E. coli* presents a bactericidal effect (Blum et al., 1992; Jung & Ahn, 2022). Considering that, three families of proteins were selected for functional characterization, HU, ClpP, and DnaJ. To select the most promising protein sequences from the total sequences retrieved by the HHM approach, we evaluated if the chosen sequences were non-truncated when compared to homologous protein sequences using BLASTp. As a result, 24 protein sequences were selected (15 for HU, 6 for ClpP, and 3 for DnaJ), with protein identity percentage of 50-85% for HU, 83-97% for ClpP, and 55-93% for DnaJ **(Table 1)**.

**Table 1.** Summary of the main characteristics of the selected proteins.

| Protein name | Source | Length (aa) | Identity percentage | Closest related microorganism* |
| --- | --- | --- | --- | --- |
| HU.d1 | Dourados | 91 | 76% | *Thermoleophilia bacterium* |
| HU.d2 | Dourados | 88 | 74% | *Solirubrobacterales bacterium* |
| HU.jz | Juazeiro | 90 | 60,23% | *Sphingomonas koreensis* |
| HU.go | mgpGO | 90 | 70,79% | Unclassified bacterium |
| ClpP.da | Atacama Desert | 198 | 97,92% | *Rubrobacteraceae bacterium* |
| ClpP.jz | Juazeiro | 193 | 83,87% | *Candidatus Saccharibacteria bacterium* |
| ClpP.z4 | Zoo 4 | 203 | 93,88% | *Actinobacteria bacterium* |
| DnaJ.tt | Tietê | 375 | 55,65% | *Rhizobium sp. ARZ01* |
| DnaJ.z4 | Zoo 4 | 389 | 88,11% | Unclassified bacterium |
| DnaJ.jz | Juazeiro | 379 | 93,90% | *Thermomonas aquatica* |

\* Considering the closest similar protein according to BLASTp hits.

### Sequence analysis of selected protein candidates

The 24 selected amino acid sequences were used to construct dendrograms for each protein family. This analysis allowed us to categorize the sequences according to protein sequence identity and select them based on their position resulting from the dendrogram built (**Figure S2-S4**). For this, we took into account as selection criteria (i) sequences presenting a lower percentage of amino acid identity; (ii) that belong to different environmental sources, and; (iii) that were located in different branches of the dendrogram. Those observations enabled us to choose ten sequences for further functional characterization (**Table 1**). To verify the presence of conserved residues, sequence alignments (**Figure S5)** and 3D structure models were constructed (**Figure S6**). In the HU sequence alignment, the presence of proline 63 in HU sequences was verified (**Figure S5A**), a highly conserved residue at the ends of the beta strands that surround the DNA and which, as part of an Arg-Asn-Pro motif (RNP), stabilizes the twists of the DNA (Grove, 2011). Interestingly, HU.d1 and HU.d2 have valine instead of arginine at position 61. The presence of R does not contribute to the stabilization of DNA twists, however, in *Thermotoga maritima*, this motif contains V instead of R, therefore its removal appears to be required to maintain any distinction between perfect duplexes and damaged DNA (Grove, 2011). Ligation to perfect duplexes DNA would allow HU protein to protect them before exposure to stress, thus reducing damage. Additionally, the conserved glycine at position 15 was also observed, a residue that mediates loop flexibility and promotes the thermal stability of HU (Georgoulis et al., 2020; Kawamura et al., 1996) (**Figure S5A**). Likewise, the presence of the Ser-His-Asp catalytic triad was observed in ClpP sequences, a motif fully conserved in members of the ClpP family and which plays an essential role in the cleavage capacity of proteases (Khan et al., 2022) (**Figure S5B**). Similarly, the conserved His-Pro-Asp motif in DnaJ sequences was verified (**Figure S5C)**. This conserved motif recognizes and stimulates the ATPase activity of DnaK (Ahmad et al., 2011). We observed that the percentage of identity ranged from 60 to more than 90% in the ten selected sequences when compared to proteins from BLASTp hits (**Table 1**), something that in principle would indicate that there is no novelty in the sequences. However, it should be noticed that none of these sequences have been experimentally characterized in previous studies.

Next, we carried out the cloning of the ten selected genes in individual genetic circuits (**Table S5**). For this, each nucleotide sequence was cloned under control of the constitutive medium expression promoter Pj*106* and the RBS BBa_B0034 (Mahajan, 2003), which was reported to present medium translation strength. Constructions were assembled into the pUC19 vector and resulting plasmids were transformed into *E. coli* DH10B (**Figure 1A**).

### Metabolic burden evaluation of the genetic constructions

Considering that metabolic load, defined as the proportion of a host cell’s resources – energy molecules or carbon building blocks – that are used to express heterologous proteins have an important impact in normal cellular processes (Glick, 1995; Wu et al., 2016), we proceeded to evaluate the metabolic burden of the ten genetic circuits by measuring the growth curves of the clones in a culture medium without any stress factor. For this, growth curves of the *E. coli* DH10B clones bearing empty plasmid or genetic constructions were made in a minimal medium over time, and the fitness cost was calculated considering the growth rates (**Figure 2A; Table S6**). Unexpectedly, the clone carrying the empty pUC19 plasmid showed the highest fitness cost (42.1%). In accordance with this observation, there are several reports found in the literature or in research discussion forums that describe difficulties related to the growth or transformation efficiency of *E. coli* cells transformed with empty plasmid pUC19 (Amirie et al., 2014; Toh, 2013). This is probably due to a higher number of copies of this plasmid when it is not bearing extra genes, which imposes a high metabolic cost on the cell. To circumvent this limitation, we perform the experiments with clones (bearing empty plasmid or genetic circuits) situated in the same metabolic state, that is, stress tests were just initiated when cultures reached the early exponential growth phase (OD_600_ = 0.3 - 0.4). Also, for comparison purposes, untransformed *E. coli* cells were used as a negative control.

**Figure 2.**
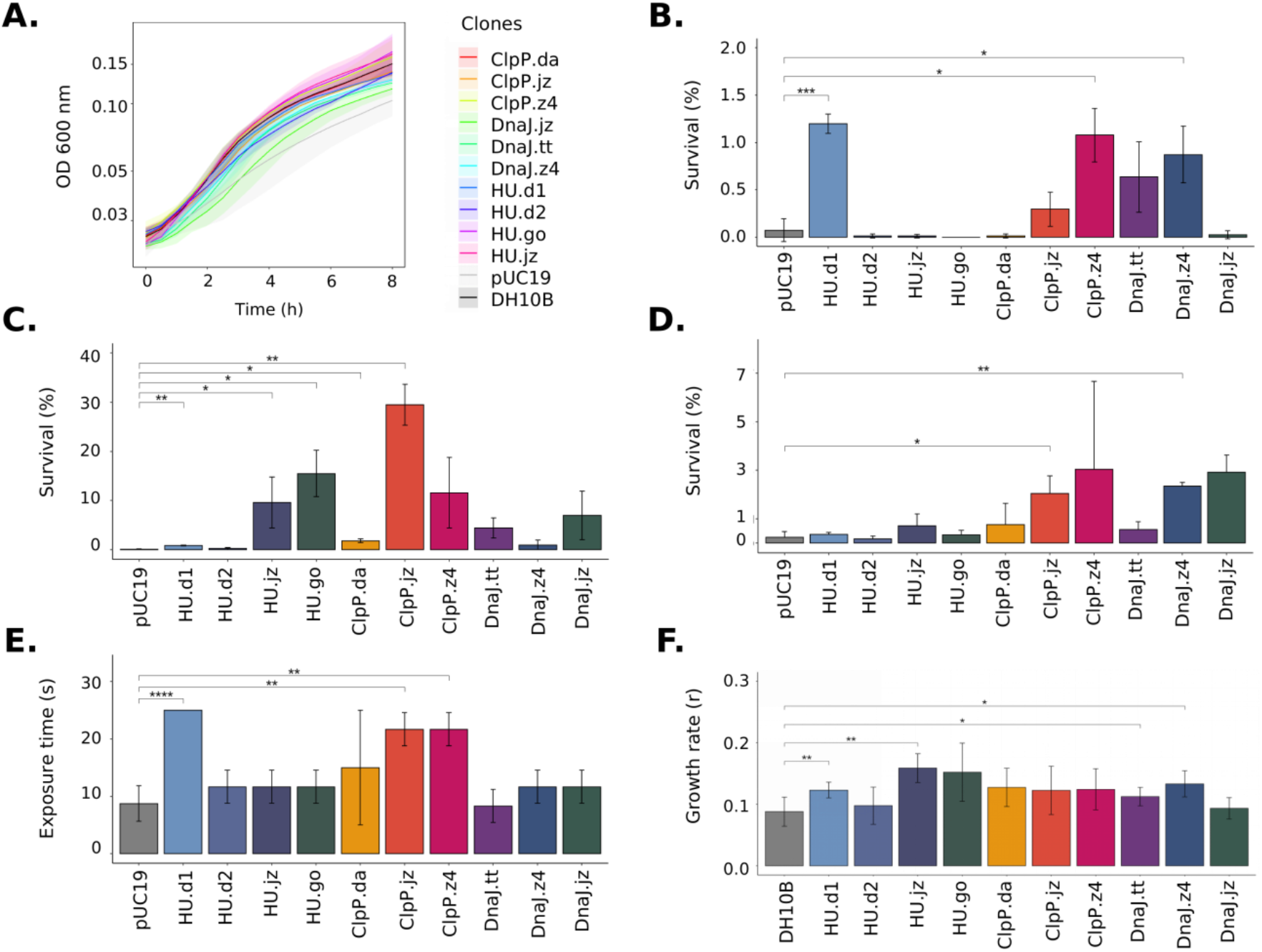
Functional characterization of *E. coli* DH10B clones carrying *hup, clpP*, and *dnaJ* genes. (A) Growth curve of *E. coli* DH10B clones in liquid M9-gly medium during 8 h; optical density at 600 nm was measured every 30 min. *E. coli DH10B* carrying empty pUC19 plasmid was used as a negative control. To test the effects of stress factors, *E. coli* DH10B clones were grown to exponential phase at 30 °C, then (B) treated at 55 °C for 30 min, (C) exposed to acid medium (pH = 3.0) for 60 min or (D) to 12 mM hydrogen peroxide for 60 min. Serial dilutions were made and CFU counts were used to determine the survival percentage. (E) Effect of UV radiation resistance in *E. coli* clones. Overnight cultures of *E. coli* clones were irradiated with a germicidal lamp (254 nm UV) during 0, 5, 10, 15, 20, and 25 s. Reaching a certain level of exposure was determined if there were at least 10 colonies in the plate. (F) Effect of salinity on growth rates of *E. coli* DH10B clones. Growth rates were determined from the exponential curve fitting function as described in Materials and Methods. Student’s t-test was used for comparison of the values obtained in the experiments and significance determination (p > 0.05; *p ≤ 0.05; **p ≤ 0.01; ***p ≤ 0.001; ****p ≤ 0.0001)

### Response to stress conditions conferred by the stress-related genes

Microorganisms resistant to several stresses play an essential role in industrial bioprocesses because they permit increased yields and lower production costs (Thorwall et al., 2020; L. Wang et al., 2022). Production process generally involves osmotic (e.g., processes involving high salt concentration culture medium), high temperature (such as processes that enable non-sterilized fermentation), acidity (e.g., organic acid production), and oxidative stresses (e.g. accumulation of radical oxygen species) (Auesukaree, 2017; Q. Li et al., 2011; Patel et al., 2006; Qin et al., 2009; Sanlier et al., 2019; Ye & Chen, 2021). Similarly, bioremediation of polluted environments is also affected by different abiotic parameters since wastewater is produced by various industries such as chemical, petrochemical, and textile (Aliko et al., 2022; Koshlaf & Ball, 2017; Le Borgne et al., 2008; Wernick et al., 2016). Another critical application of stress-resistance microorganisms is in enzyme production processes, in which bacterial hosts need to be active and stable at high salt contents or high temperatures (Margesin & Schinner, 2001; Verma et al., 2021).

In order to study if synthetic genetic circuits – bearing independent new sequences coding for the DNA-binding protein HU, the ATP-dependent Clp protease proteolytic subunit, and the molecular chaperone DnaJ – increase *E. coli* resistance to diverse stress conditions, we perform stress shocks assays with *E. coli* clones in five stress conditions, including high temperature, acidity, oxidative and osmotic stress, and UV radiation (**Figure 1A; Table S7**).

Considering the set of assays in the five stress conditions, we observed that expression of most of the encoding sequences granted stress resistance to the cells in differing degrees (**Figure 2**). Cells expressing *hu*.*d1* showed a positive response to four stress factors over the ones expressing either of the other genes, with an average survival percentage of over 1.2% in heat shock tests (**Figure 2B**) and 0.8% in the acidic challenge (**Figure 2C**). These values represented a fold change of 17 and 9, respectively, in comparison to *E. coli* cells harboring the empty plasmid (0.07% in heat shock assays and 0.1% in acid shock experiments). These results agree with the previous work of Ogata and co-workers (1997), which showed that the HU protein maintains the negative supercoiling of DNA during thermal stress and contributes to cellular thermotolerance in *E. coli* (Ogata et al., 1997). Similarly, the HU role in acid resistance was also corroborated by several studies (Alves et al., 2023; Bi et al., 2009; de Siqueira et al., 2020; Guazzaroni et al., 2013). In the UV radiation tests, cells expressing *hu*.*d1* also showed a higher percentage of survival (colony growth until 25 s of UV radiation exposure) compared with control cells (growth until 10 s) (**Figure 2E**). This is probably because HU acts against UV radiation stress by preventing the formation of photoproducts and in the RecA-dependent repair process (Miyabe, 2000). At the same time, in the salinity assays, cells expressing *hu*.*d1* reached a growth rate of 0.12. This rate was 33% higher than that achieved by untransformed *E. coli* cells (**Figure 2F**). The HU protein response to osmotic stress would involve the modulation of expression of sigma factor *rpoS* (**σ**38), a global regulator whose translation is stimulated in response to high osmolarity (Battesti et al., 2011; Muffler et al., 1996).

In a similar way, cells expressing *clpP*.*jz* or *dnaJ*.*z4* related sequences showed a positive response when exposed to three different stress factors (acidity, oxidative stress, and UV radiation for *clpP*.*jz*, and heat shock, oxidative, and osmotic stress for *dnaJ*.*z4*). For example, cells expressing *clpP*.*jz* showed an average survival percentage of around 29% in the acidity challenge (**Figure 2C**) and 2% in oxidative stress (**Figure 2D**). These values represent a fold change of 290 and 9 times, respectively, in comparison to negative controls. Moreover, UV radiation tests also showed a significantly higher survival of cells expressing *clpP*.*jz* compared with cells bearing empty plasmid (20 s versus 10 s) (**Figure 2E**). In accordance with the obtained results, there are several works in the literature describing a connection between this protease and cellular damage protection in stress conditions (L. Chen et al., 2015; de Siqueira et al., 2020; Guazzaroni et al., 2013). For instance, a functional metagenomic study from an extremely acidic environment identified a *clpP* gene able to confer resistance to acidity and UV radiation in *E. coli* (Guazzaroni et al., 2013). Also, when this gene was used for the construction of a combinatorial acid stress-tolerance module improved the growth robustness of *E. coli* at low pH (de Siqueira et al., 2020). Additional examples in other microorganisms, such as *Bifidobacterium longum* and *Lactococcus lactis*, also show that ClpP is involved in the acid stress response (Frees & Ingmer, 1999; Wei et al., 2015). It is important to highlight that during oxidative stress, hydrogen peroxide is easily diffused into cells, increasing intramolecular concentrations (Sen et al., 2020). In the cytoplasm, the reactions generate hydroxyl radicals that damage DNA and other biomolecules – similar to the presence of protons during acid stress –, endangering the survival of the cell (Sen et al., 2020; **Figure 3**). In general, hydrogen peroxide does not cause protein aggregation, however, the oxidation of proteins by reactive oxygen species (ROS) requires misfolded-protein degradation (Daly et al., 2007; S. Wang et al., 2009). Similarly, after UV radiation exposure, excited sensitizers can directly react with DNA directly by transferring electrons or energy to molecular oxygen, thus generating ROS, which in turn can damage the DNA molecule (Schuch & Menck, 2010).

**Figure 3.**
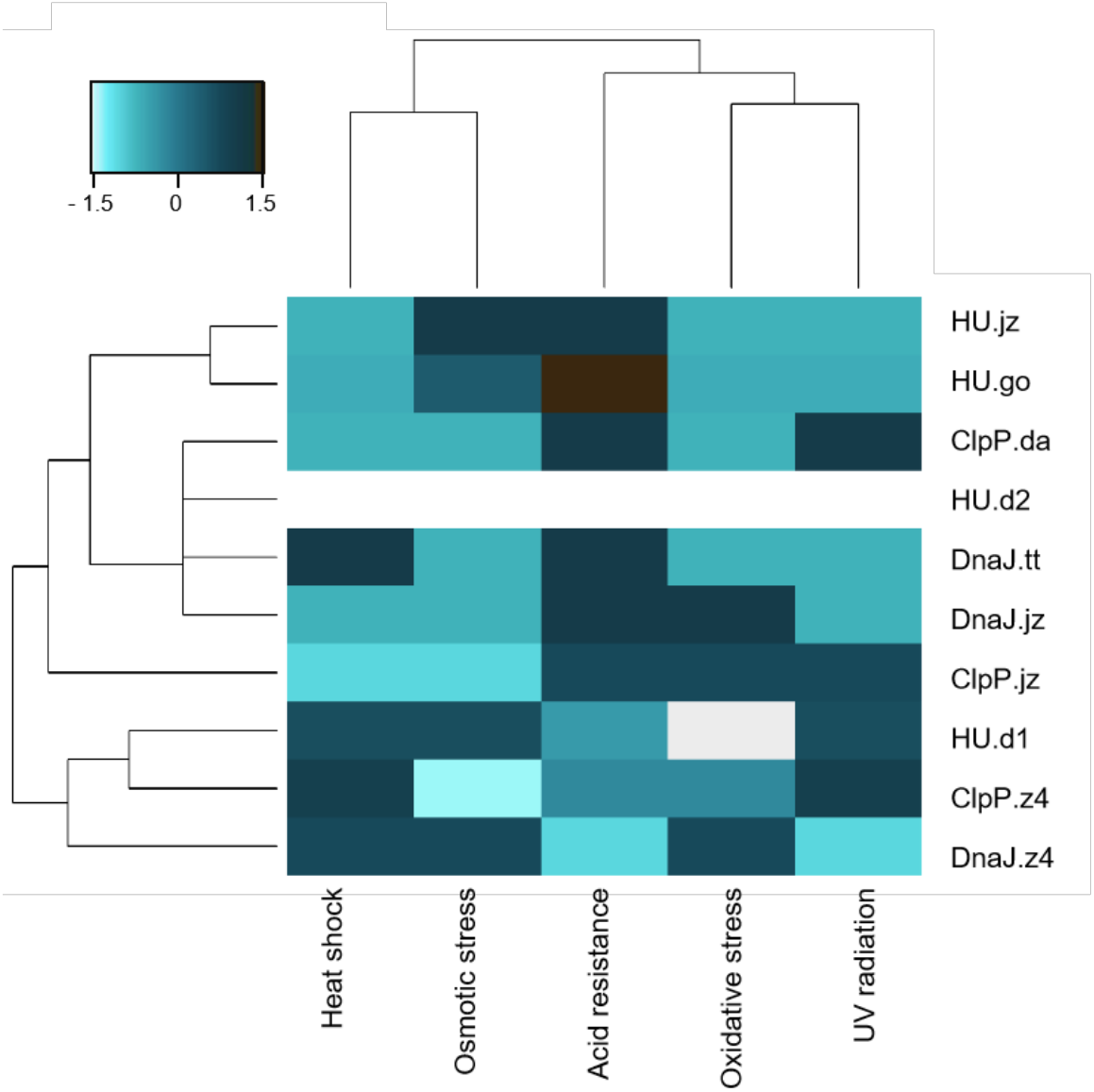
Clustering of *E. coli* clones considering functional characterization in the five stress conditions. Hierarchical cluster of *E. coli* DH10B clones carrying the pUC19 plasmid with genes *clpP, hup*, and *dnaJ*. Scale bar denotes the Z-score fold change. Strong color represents an increased response level to stress factors.

On the other hand, clones expressing *dnaJ*.*z4* had an average survival of 0.87% in the heat shock test (**Figure 2B**) and 2.35% in oxidative stress (**Figure 2D**), representing more than 12 and 10 times of survival in respective negative control. Also, when the cells expressing *dnaJ*.*z4* were subjected to saline stress, they reached a 44% higher growth rate compared to untransformed *E. coli* cells (**Figure 2F**). It should be noticed that the DnaJ protein is part of the molecular response machinery to heat shock, confirming the results found in our study (Ghafoori et al., 2017). A recent work in which the *dnaJ* gene from *Bacillus halodurans* was overexpressed in *E. coli* showed improved cell thermotolerance, indicating that DnaJ overexpression was sufficient to increase resistance to high temperatures (Ghafoori et al., 2017). On the other hand, our study is the first one – to the best of our knowledge – to report DnaJ protein protection against oxidative cellular damage.

## Conclusions

The approach presented here allowed us to identify five novel coding sequences conferring resistance to at least two stress conditions when expressed in *E. coli*. It should be noticed that all expressed proteins, except for HU.d2, demonstrated resistance to at least one of the tested stress conditions (**Figure 2**; **Figure 3**). Notably, among all synthetic genetic constructions, cells expressing HU.d2 exhibited a significantly higher metabolic cost compared to the other clones (**Figure 2A**; **Table S6**), which should explain its limited response to stress conditions. Also, it is worth mentioning that the response level varied among clones despite proteins belonging to the same family. This inconsistency may be attributed to the origin of metagenomic DNA from diverse microorganisms, where certain genes tend to be preferentially expressed in hosts possessing transcription/translation machinery more closely resembling the source organism (Gabor et al., 2004).

Clustering of *E. coli* clones stress-responses illustrated that probably the expression of each protein corresponds to specific molecular mechanisms of survival (**Figure 3**). Notably, UV radiation, acidity, and oxidative stress exhibit a similar response pattern, probably due to the fact that UV radiation and acidity indirectly induce oxidative stress by generating ROS (Schuch & Menck, 2010).

Despite considerable research on stress response available in the literature, the approach presented here has revealed novel aspects of this well-established response, including genes that potentially enhance resistance. On the other hand, from a biotechnological point of view, one of the identified novel coding sequences (*hu*.*d1*) retrieved from an acidic soil metagenome notably increased microbial tolerance to four different stress conditions, indicating its suitability for the construction of a synthetic circuit directed to expand broad bacterial resistance. Moreover, the assembly of genetic circuits with a combination of the best-identified gene candidates from our study (e.g., *hu*.*d1, clpP*.*jz*, and *dnaJ*.*z4*) should lead to improved bacterial endurance. Taken together, this work provides novel biological parts for the engineering of microbial hosts which could be used for extreme industrial applications. Still, further research is necessary to comprehend the mechanisms of action when these proteins are heterologously expressed and to understand the functional behavior of the identified genes in other hosts than *E. coli*.

## Supplemental Material

Figure S1. Overview of the pipeline for analysis and identification of potential stress-related genes.

Figure S2. Dendrogram of the HU protein sequences.

Figure S3. Dendrogram of the ClpP protein sequences.

Figure S4. Dendrogram of the DnaJ protein sequences.

Figure S5. Multiple sequence alignment of HU, ClpP, and DnaJ protein sequences.

Figure S6. 3D structural model of selected proteins.

Figure S7. Survival profiles of clones harboring *hup, dnaJ*, and *clpP* genes.

Figure S8. Effect of the retrieved genes in UV radiation resistance.

Table S1. Description of metagenomes selected through the MG-RAST platform.

Table S2. Assembly statistics of metagenomic datasets by MegaHit software.

Table S3. Summary of annotation results made by the Prokka software.

Table S4. Summary of the number of hits of different proteins found in metagenomes.

Table S5. Amino acid sequences of the selected proteins for experimental validation

Table S6. Growth rate and fitness cost of clones grown in minimal medium.

Table S7. Summary of the results obtained for each stress assay.

## Author Contributions

MEG and RSR designed the project. JHJ and ENS performed experiments and *in silico* analysis. NHS developed the HMM profiles and metagenome mining. JHJ and MEG wrote the manuscript. All authors critically revised the manuscript and approved the submitted version.

## Acknowledgments

The authors would like to thank the lab technician Thalita Riul Prado for her invaluable assistance in the course of this work.

