## Supplemental Figures S1-S8 and Table S1-S7 for "Identification and functional analysis of novel stress-resistance genes from metagenomes of extreme environments"

### Supplementary material

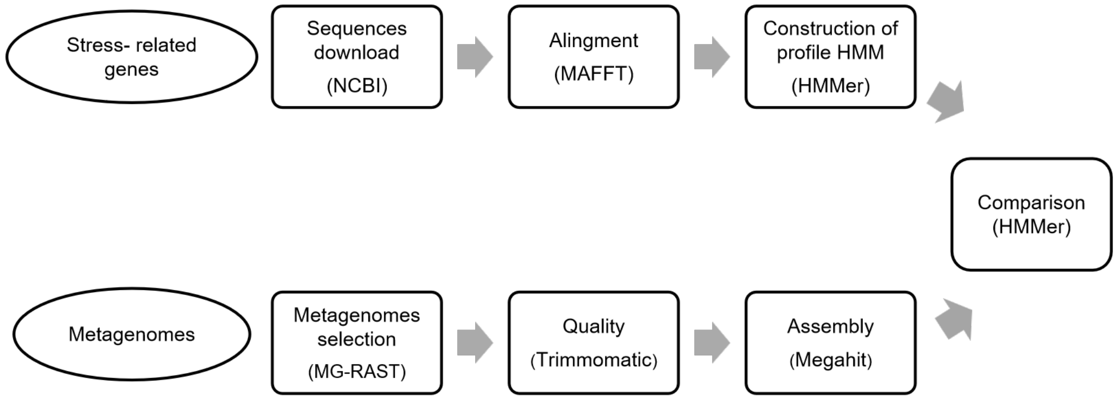


**Figure S1.** Overview of the pipeline for analysis and identification of potential stress-related genes.

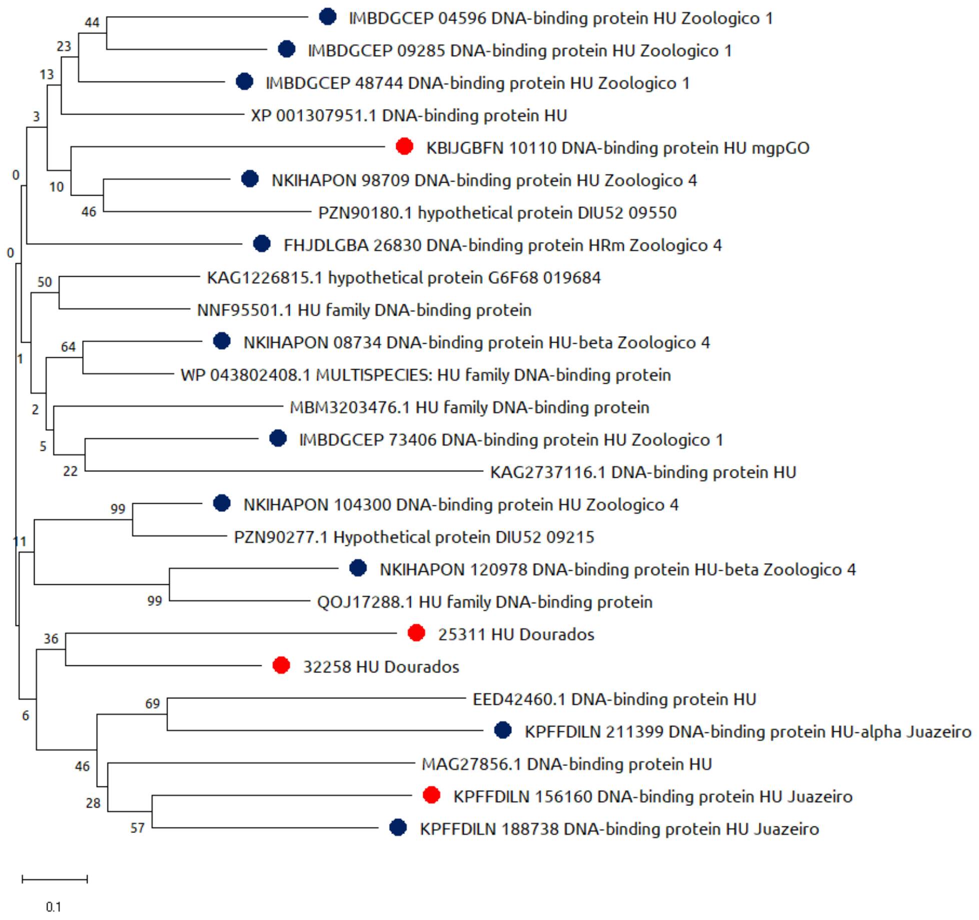


**Figure S2.** Dendrogram of the HU family. Dendrogram of 26 HU protein sequences using the Neighbor-Joining (NJ) method. Amino acid sequences were retrieved from GenBank as indicated by accession numbers as follows: XP_001307951.1 (*Trichomonas vaginalis* G3), MBM3203476.1 (*Candidatus Woesearchaeota archaeon*), KAG2737116.1 (*Suillus brevipes* Sb2), PZN90180.1 (bacteria), QOJ17288.1 (*Phycisphaeraceae bacteria*), PZN90277.1 (bacteria), WP_0438.1 (*Arenimonas*), NNF95501.1 (*Halobacteria archaeon*), KAG1226815.1 (*Rhizopus microsporus*), EED42460.1 (*Enterocytozoon bieneusi* H348), MAG27856. 1 (*Candidatus Pacearchaeota archaeon*). The numbers on the nodes correspond to the bootstrap percentage values for 1000 replicates. The circles represent the pre-selected sequences, and the color if they were selected (red) or not (blue).

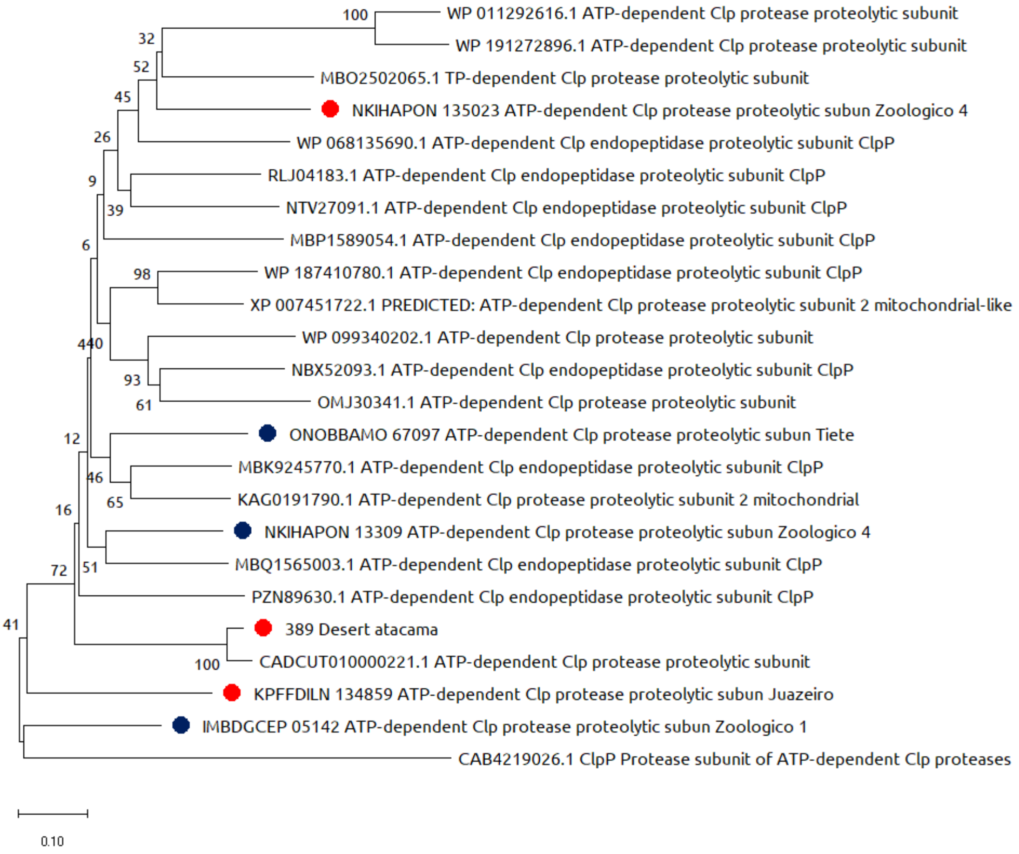


**Figure S3.** Dendrogram of the ClpP family. Dendrogram of 24 ClpP protein sequences using the Neighbor-Joining (NJ) method. Amino acid sequences were retrieved from GenBank as indicated by accession numbers as follows: WP_011292616.1 (*Thermobifida fusca*), WP_191272896.1 (*Nocardiopsis terrae*), MBO2502065.1 (*Thermoanaerobacterales bacterium*), WP_068135690.1 (*Limnochorda pilosa*), RLJ04183 .1 (*Candidatus Aenigmarchaeota archaeon*), NTV27091.1 (*Methanothrix* sp.), MBP1589054.1 (*Kiritimatiellae bacterium*), WP_187410780.1 (*Saccharophagus* sp. K07), XP_007451722.1 (*Lipotes vexillifer*), WP_099340202.1 (*Candidautus Fonsibacter*), NBX52093.1 (*Proteobacteria bacterium*), OMJ30341.1 (*Smittium culicis*), MBK9245770.1 (*Burkholderiales bacterium*), KAG0191790.1 (*Apophysomyces* sp. BC1034), MBQ1565003.1 (*Clostridia bacterium*), PZN89630.1 (bacterium), CADCUT010000221.1 (uncultured *Rubrobacteraceae bacterium*), CAB4219026.1 (uncultured *Caudovirales* phage). The numbers on the nodes correspond to the bootstrap percentage values for 1000 replicates. The circles represent the pre-selected sequences, and the color, if they were selected (red) or not (blue).

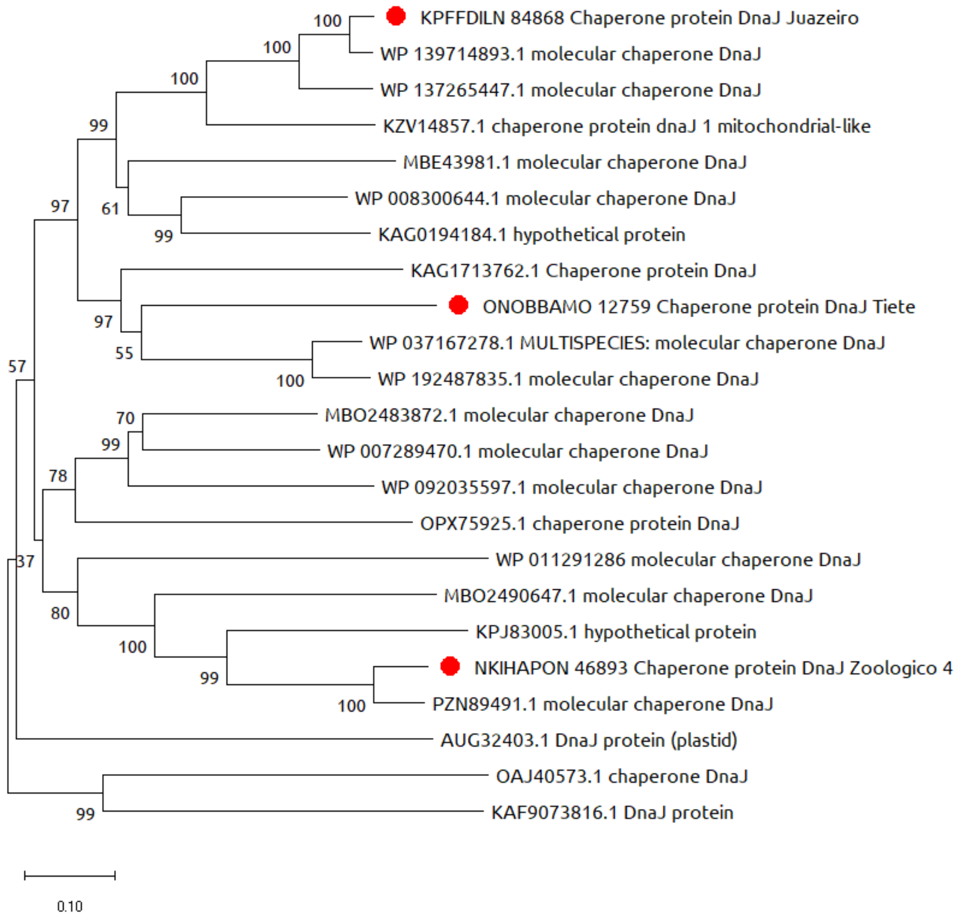


**Figure S4.** Dendrogram of the DnaJ family. Dendrogram of 23 amino acid sequences of the DnaJ protein using the Neighbor-Joining (NJ) method. Amino acid sequences were retrieved from GenBank as indicated by accession numbers as follows: WP_139714893.1 (*Thermomonas* sp. SY21), WP_137265447.1 (*Luteimonas gilva*), KZV14857.1 (*Dorcoceras hygrometricum*), MBE43981.1 (*Thaumarchaeota archaeon*) , WP_008300644.1 (*Candidatus Nitrosopumilus salaria*), KAG0194184.1 (*Apophysomyces* sp. BC1034), KAG1713762.1 (*Nymphon striatum*), WP_037167278.1 (unclassified *Rhizobium*), WP_192487835.1 (*Agrobacterium* sp. AGB081), MBO481 *Firmicutes bacterium*), WP_007289470.1 (*Thermosinus carboxydivorans*), WP_092035597.1 (*Planifilum fulgidum*), OPX75925.1 (*Methanosaeta* sp. PtaB.Bin018), WP_011291286 (*Thermobifida fusca*), MBO2490647.1 (*Rhodo Kthermus*J5P8.1) *Gemmatimonas* sp.SG8_23), PZN89491.1 (bacterium), AUG32403.1 (*Paulinella longichromatophora*), OAJ40573.1 (*Batrachochytrium dendrobatidis* JEL423), KAF9073816.1 (*Rhodocollybia butyracea*). The numbers on the nodes correspond to the bootstrap percentage values for 1000 replicates. The circles represent the pre-selected sequences, and the color, if they were selected (red) or not (blue).

A.

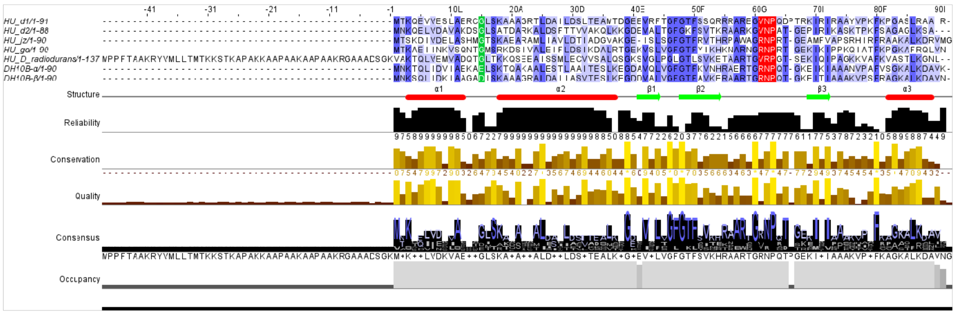

B.

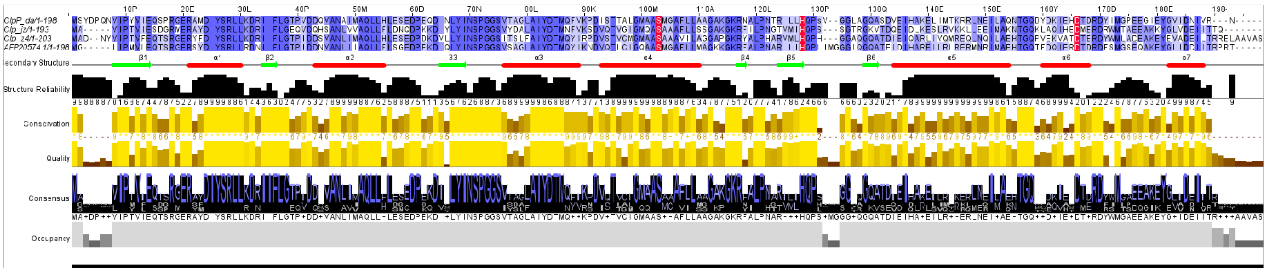

C.

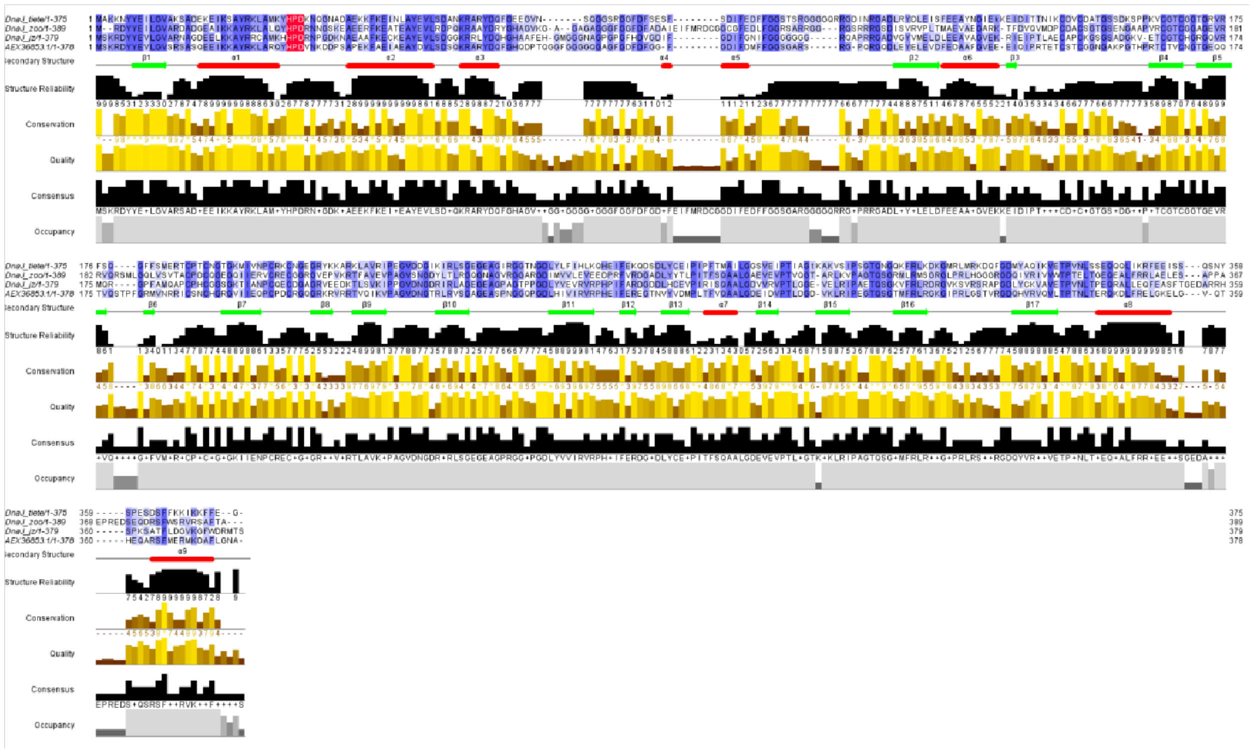

**Figure S5.** Multiple sequence alignment of HU, ClpP and DnaJ sequences. (A) Alignment of HU protein sequences, including the 4 selected sequences (HU.d1, HU.d2, HU.jz and HU.go), HU alpha and beta from *E. coli* DH10B and HU from *Deinococcus radiodurans*. (B) Alignment of ClpP protein sequences, including the 3 selected sequences (ClpP.da, ClpP.jz, and ClpP.z4) and ClpP of metagenomic origin (GenBank: JX219770). (C) Alignment of DnaJ protein sequences, including the 3 selected sequences (DnaJ.tt, DnaJ.z4, and DnaJ.jz) and DnaJ from *Alicyclobacillus acidoterrestris*. Residues from the catalytic site of each protein are indicated in red. Secondary structures and predictive reliability of proteins are shown below the sequence.

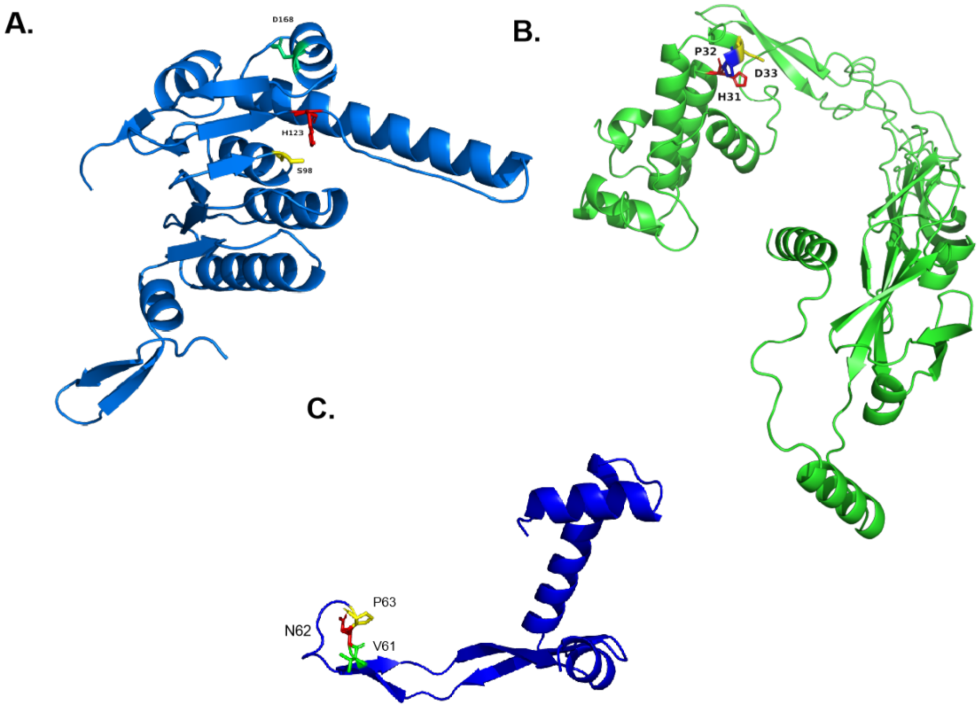


**Figure S6.** 3D structural model of selected proteins. The figure shows models for (A) DnaJ.z4, (B) ClpP.jz, and (C) HU.d1. Catalytic residues are highlighted for each protein.

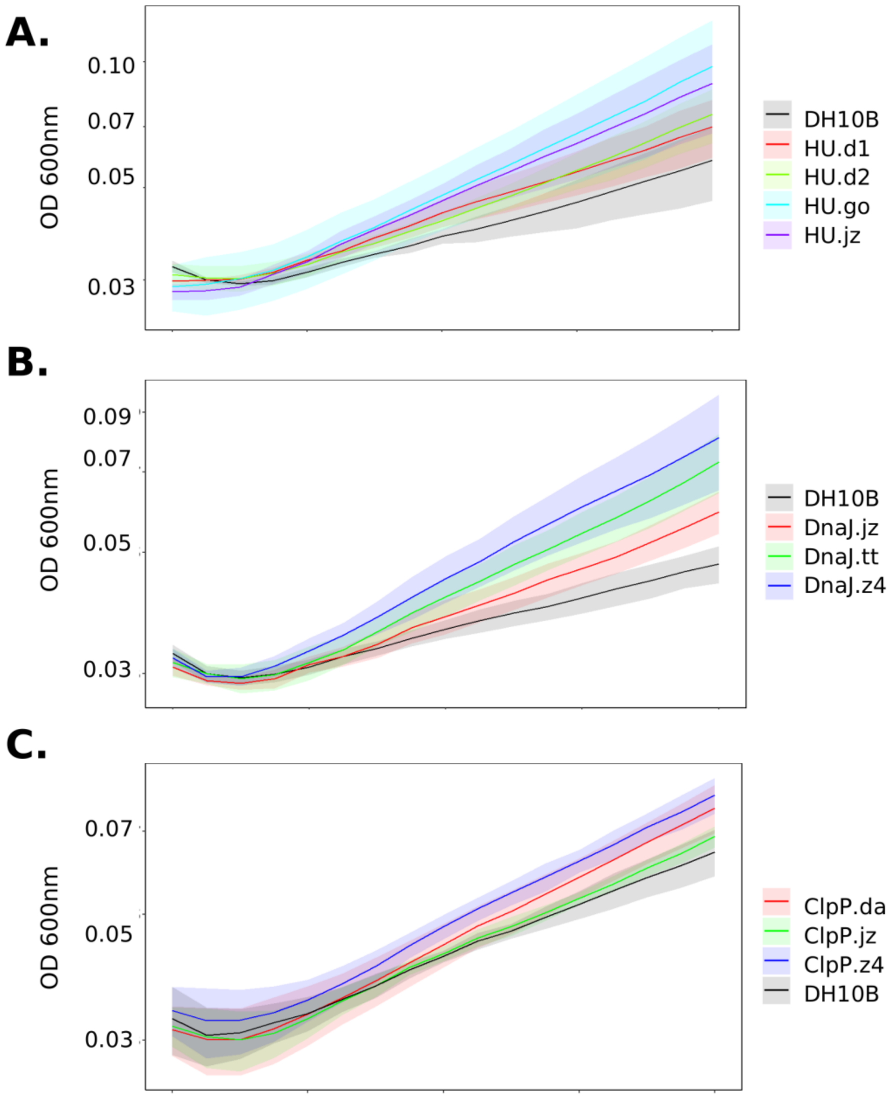


**Figure S7.** Survival growth curves of clones harboring plasmids containing (A) *hu*, (B) *dnaJ* and (C) *clpP* genes at 3.5% NaCl supplemented M9 medium.

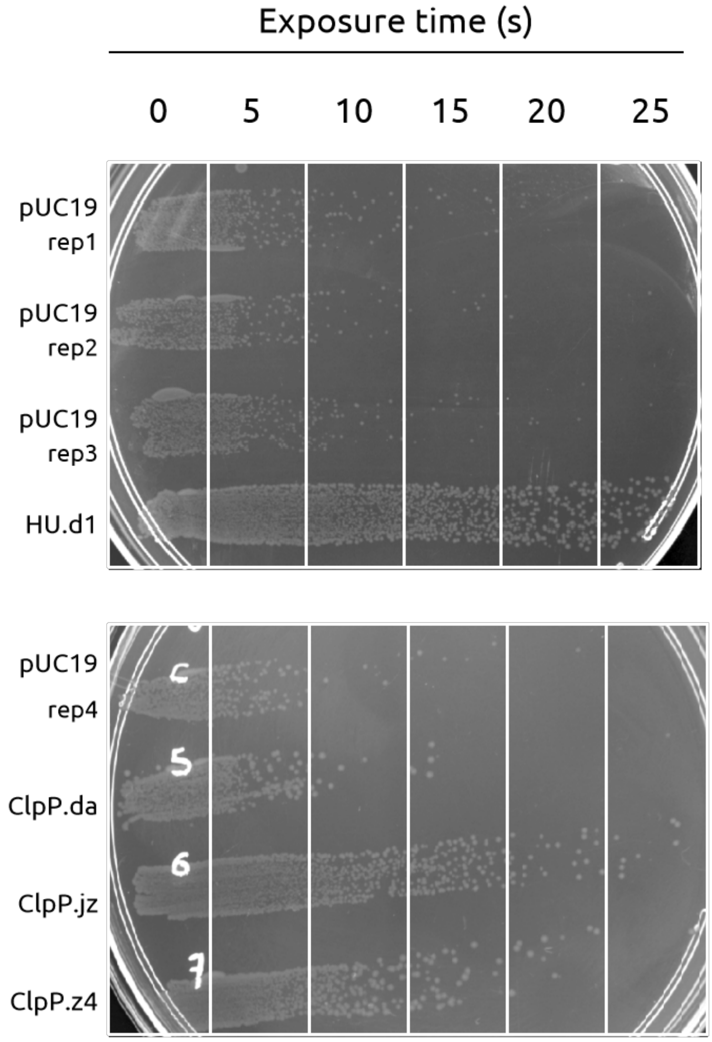


**Figure S8.** Effect of the retrieved genes in UV radiation resistance assays. Overnight cultures of *E. coli* DH10B, linearly spread on M9 plates, carrying the pUC19 plasmid with subcloned genes *clpP*, *hu* and *dnaJ* were irradiated with a germicidal lamp during 0, 5, 10, 15, 20, and 25 seconds. *E. coli* DH10B carrying empty pUC19 was used as a negative control. Each assay was performed at least three times using independent cultures.

**Table S1.** Description of metagenomes selected through the MG-RAST platform.

| ID | File name | Study name | Size (MB) | Material | Condition | Location |
|---|---|---|---|---|---|---|
| mgp8766 | AmazPluma | AmazPluma | 1300 | River water | Salinity | Amazon, Brazil |
| mgp81659 | juazeiro | Microbial community urban environment with heavy metal industrial waste | 299.3 | Margem do rio | High heavy metals concentration levels (Zn and Cu) | Ceara, Brazil |
| mgp18106 | mgpGO | Evaluation of microbiota in serpentine soils for nickel biomining | 1400 | Bulk soil | High heavy metals concentration levels (Ni) | Goias, Brazil |
| mgp86353 | Tiete | Plas_Tiete | 10400 | Water | Contaminated river | São Paulo, Brazil |
| mgp5435 | Zoologico 1 | Study of composting at the São Paulo Zoo (1) | 368.7 | Compost | Temperature 66-67º C | São Paulo, Brazil |
| mgp5435 | Zoologico 2 | Study of composting at the São Paulo Zoo (2) | 302.6 | Compost | Temperature 66-67º C | São Paulo, Brazil |
| mgp5435 | Zoologico 3 | Study of composting at the São Paulo Zoo (3) | 1500 | Compost | Temperature 66-67º C | São Paulo, Brazil |
| mgp5435 | Zoologico 4 | Study of composting at the São Paulo Zoo (4) | 1900 | Compost | Temperature 66-67º C | São Paulo, Brazil |
| mgp11367 | Desert_atacama | Atacama metagenomes | 1.2 | Rock | High temperature | Chile |
| mgp18410 | Dourados | Metagenomas solos de Dourados | 46.7 | Soil | Acidity | Mato Grosso, Brazil |

**Table S2.** Assembly statistics of metagenomic datasets by MegaHit software.

| Study place | Time (h:m:s) | File size | Total length (bp) | N50 (bp) | Total contigs | Min (bp) | Max (bp) | Mean (bp) |
|---|---|---|---|---|---|---|---|---|
| Amazon | 1:05:05 | 166.4 M | 153839154 | 509 | 300170 | 200 | 105781 | 512 |
| Juazeiro | 0:36:37 | 305.5 M | 277741274 | 399 | 664071 | 201 | 10585 | 418 |
| Barro Alto | 01:09:49 | 40.1 M | 36771520 | 484 | 81125 | 200 | 3313 | 453 |
| Tietê | 02:12:23 | 190.9 M | 178760935 | 635 | 290434 | 211 | 117691 | 615 |
| Zoo 1 | 00:15:05 | 106.8 M | 98014883 | 426 | 212382 | 200 | 21393 | 461 |
| Zoo 2 | 0:12:57 | 94.6 M | 86253073 | 404 | 201129 | 200 | 8025 | 428 |
| Zoo 3 | 00:38:03 | 122.8 M | 113058559 | 432 | 234957 | 200 | 57509 | 481 |
| Zoo 4 | 1:10:48 | 272.4 M | 250706729 | 452 | 519263 | 200 | 37665 | 482 |

**Table S3.** Summary of the annotation results made by the Prokka software.

| Study place | Annotation | CDS | Hypothetical proteins | Non hypothetical proteins |
|---|---|---|---|---|
| Amazon | 238677 | 237951 | 228208 | 9743 |
| Juazeiro | 214382 | 211222 | 165074 | 46148 |
| Barro Alto | 46982 | 46289 | 41059 | 5230 |
| Tietê | 155070 | 152095 | 113505 | 38590 |
| Zoo 1 | 77408 | 75542 | 60420 | 15122 |
| Zoo 2 | 65510 | 63935 | 49206 | 14729 |
| Zoo 3 | 91203 | 89077 | 65406 | 23671 |
| Zoo 4 | 203237 | 199648 | 142173 | 57475 |

**Table S4.** Summary of the number of hits of different proteins found in the metagenomes.

| Protein | M1 | M2 | M3 | M4 | M5 | M6 | M7 | M8 | M9 | M10 |
|---|---|---|---|---|---|---|---|---|---|---|
| **DPS** | 4 | 0 | 3 | 32 | 2 | 5 | 5 | 33 | * | 4 |
| **HU** | 18 | 98 | 15 | 84 | 36 | 38 | 43 | 93 | * | 15 |
| **RBP** | 1 | 28 | 4 | 29 | 7 | 6 | 6 | 22 | * | 5 |
| **DnaA** | 60 | 62 | 6 | 85 | 30 | 28 | 45 | 97 | 1 | 23 |
| **GyrA** | 31 | 35 | 9 | 64 | 33 | 20 | 25 | 74 | 1 | 29 |
| **RecA** | 186 | 22 | 5 | 86 | 24 | 12 | 26 | 54 | * | 24 |
| **ClpA** | 99 | 84 | 27 | 206 | 64 | 54 | 106 | 186 | 1 | 111 |
| **ClpC** | 84 | 78 | 25 | 183 | 63 | 42 | 94 | 177 | 1 | 103 |
| **ClpE** | 60 | 62 | 18 | 161 | 62 | 37 | 79 | 152 | 1 | 67 |
| **ClpL** | 50 | 45 | 19 | 135 | 42 | 28 | 62 | 114 | 1 | 58 |
| **ClpP** | 68 | 54 | 11 | 101 | 28 | 26 | 43 | 87 | 1 | 17 |
| **ClpX** | 76 | 64 | 18 | 167 | 47 | 34 | 72 | 143 | * | 71 |
| **Cas1** | 2 | 5 | 0 | 0 | 1 | 5 | 7 | 12 | * | * |
| **Cas2** | 0 | 1 | 0 | 0 | 0 | 5 | 7 | 11 | * | * |
| **Cas9** | 5 | 3 | 1 | 8 | 1 | 3 | 2 | 6 | * | * |
| **DnaJ** | 22 | 90 | 9 | 114 | 42 | 23 | 33 | 105 | * | * |
| **DnaK** | 36 | 45 | 3 | 81 | 20 | 27 | 45 | 58 | * | * |

The columns represent the following environments: (M1) Amazon River (M2) Juazeiro (M3) Serpentine soil (M4) Tietê River (M5) Zoo 1 (M6) Zoo 2 (M7) Zoo 3 (M8) Zoo 4 (M9) Desert from Atacama (M10) Dourados.

**Table S5.** Amino acid sequences of the 10 selected proteins for experimental validation.

| Sequence name | Aminoacid sequence |
|---|---|
| HU_d1 | MTKQEVVESLAERCGLSKAAAGRTLDAILDSLTEAMTDGEEVRFTGFGTF SSQRRRAREGVNPQDpTRKIRIRAAYVPKFKPGASLRAAIR |
| HU_d2 | MNKQELVDAVAKDSGLSATDARKALDSFTTVVAKQLKKGDEVALTGFG KFSVTKRAARKGVNPATGEPIRIKASKTPKFSAGAGLKSA |

| | |
|---|---|
| HU_jz | MTSKDIVDELASHMGTSKAEARAMLIAVLDTIADGVAKGEISLSGFGTFRV THRPAWAGRNPRTGEAMFVAPSRHIRFRAAKALKDRVMG |
| HU_go | MTKAEIINKVSQNTGMSRKDSIVALEIFLDSIKDALRTGEKVSLVGFGTFYI KHKNARNGRNPRTGEKIKIPPKQIATFKPGKAFRQLVN |
| ClpP_da | MSYDPQNVIPYVIEQSPRGERAMDIYSRLLKDRIIFLGTPVDDQVANAIMA QLLHLESEDPEQDINLYINSPGGSVTAGLAIYDTMQFVKPDISTTALGMAA SMGAFLLAAGAKGKRNALPNTRILLHQPsVGGLAGQASDVEIHAKELIMT KRRLNEILAQNTGQDYDKIEHDTDRDYIMGPEEGIEYGVIDNIVRN |
| ClpP_jz | MAVIPTVIESDGRMERAYDIYSRLLKDRIIFLGEQVDQHSANLVVAQLLFL DNQDPKKDIYLYINSPGGSVYDALAIYDTMNFVKSDVQTVGIGMQASAAA FLLSSGAKGKRFILPNGTVMIHQPSSGTRGKVTDQEIDLKESLRVKKLLEEI MAKNTGQKLAQIHEDMERDRWMTAEEAKKYGLVDEIITTQ |
| ClpP_z4 | MADNYVIPTVFEQTSRGERYFDIYSRLLKDRIIFLGTPIDDSVANLIMAQLL HLESEDPDKDVLLYINSPGGSITSLFAIYDTMQYIRPDVSTVCMGMAASAA AVILAGGAPGKRYALPHARVMLHQPHGGAQGQATDIEIQARLIVQMREQL NQILAEHTGQPVEKVATDTERDYWMLADEAKEYEVADEILTRRELAAVA S |
| DnaJ_tt | MAKKNYYEILGVAKSADEKEIKSAYRKLAMKYHPDKNQGNADAEKKFK EINLAYEVLSDANKRARYDQFGEEGVNSQGGSRGGFDFSESFSDIFEDFFG GSTsrgGGGQRRGDINRGADLRYDLEISFEEAYNGIEkKEIDITTNIKCDVCD ATGSSDKSPPKVCGTCGGTGRVRFSQGFFSMERTCPTCNGTGKMIVNPCR KCNGEGRYKKARKLAVRIPEGVDDGIKIRLSGEGEAGIRGGTNGDLYLFIH LKQHEIFEKQDSDLYCEIPIPFTMAILGGSVEIPTIAGtKAKVSIPSGTQNGQK FRLKDKGMRLMRKDQFGDMYAQIKVETPVNLSSEQQQLIKRFEEISSQSN YSPESDSFFKKIKKFFEG |
| DnaJ_z4 | MRDYYEILGVARDADGEAIKKAYRKLALQYHPDRNNGSKEAEERFKEAT EAYEVLRDPQKRAAYDRYGHAGVKGAGAGAGGGFGGFDFADaleIFmrdcg GCGFEDLFGGRSARRggrgSRRRGSDISVRVPLTMAEVAEGARKTFDVQVM DPCDACSGTGSENGAAPVRCGTCGGAGEVRRVQrsmlGQLVSVTACPDCQ GEGQIIERVCRECGGRGVEPVKRTFAVEVPAGVSNGDYLTLRGQGNAGVR GGARGDIMVVLEVEEDPRFVRDGADLYYTLPITFSQAALGAEVEVPTVQG TARLKVPAGTQSGRMLRMSGRGLPRLHGGGRGDQIVRIVVWTPTELTGE QEALFRRLAELESAPPAEprEDSEQDRSFWSRVRSAFTA |

| | |
|---|---|
| DnaJ_jz | MSKRDYYEVLGVARNAGDEELKKAYRRCAMKHHPDRNPGDKNAEAAF KECKEAYEVLSDGGKRRLYDQHGHAAFEHGMGGGNAGPGpgfHDVGDIF GDIFGNIFGGGggGGRQAPRRGADVGYVMELDLEEAVAGVEKRIEIPTLAE CAPCKGSGSADGKVETCGTCHGRGQVRMQRGPFAMQAPCPHCGGSGKTI ANPCQECDGAGRVEEDKTLSVKIPPGVDNGDRIRLAGEGEAGPAGTPPGD LYVEVRVRPHPIFARDGDDLHCEVPIRISQAALGDVVRVPTLGGEVELRIP AETQSGKVFRLRDRGVKSVRSRAPGDLYCKVAVETPVNLTPEQRALLEQF EASFTGEDARRHSPKSATFLDGVKGFWDRMTS |

**Table S6.** Growth rate and fitness cost of clones grown in minimal medium.

| Clone | Growth rate ($h^{-1}$) | Fitness cost (%) |
|---|---|---|
| DH10B (reference) | 0.38 | 0.0 |
| HU.go | 0.42 | -10.5 |
| HU.jz | 0.40 | -5.3 |
| ClpP.da | 0.38 | 0.0 |
| ClpP.z4 | 0.36 | 5.3 |
| DnaJ.tt | 0.36 | 5.3 |
| HU.d1 | 0.34 | 10.5 |
| DnaJ.z4 | 0.34 | 10.5 |
| ClpP.jz | 0.34 | 10.5 |
| HU.d2 | 0.28 | 26.3 |
| DnaJ.jz | 0.26 | 31.6 |
| pUC19 | 0.22 | 42.1 |

**Table S7.** Summary of the results obtained for each stress assay.

| Clone | Heat Shock (%) | Acidity (%) | Salinity ($h^{-1}$) | Oxidative stress (%) | UV radiation (s) |
|---|---|---|---|---|---|
| **Negative control** | 1.2 ± 0.10 | 0.1 ± 0.08 | 0.09 ± 0.02 | 0.24 ± 0.23 | 8.75 ± 3.1 |
| **HU.d1** | 0.01 ± 0.02 | 0.84% ± 0.12 | 0.12 ± 0.01 | 0.35 ± 0.09 | 25 |
| **HU.d2** | 0.01 ± 0.02 | 0.21 ± 0.21 | 0.10 ±0.03 | 0.18 ± 0.12 | 11.7 ± 2.9 |
| **HU.jz** | 0 | 9.60 ± 5.17 | 0.16 ± 0.02 | 0.72 ± 0.49 | 11.7 ± 2.9 |
| **HU.go** | 0.01 ± 0.02 | 15.48 ± 4.72 | 0.15 ±0.05 | 0.3 ± 0.18 | 11.7 ± 2.9 |
| **ClpP.da** | 0.3 ± 0.18 | 1.78 ± 0.38 | 0.13 ±0.03 | 0.76 ± 0.88 | 15 ± 10 |
| **ClpP.jz** | 1.08 ± 0.28 | 29.48 ± 4.19 | 0.12 ±0.04 | 2.05 ± 0.72 | 21.7 ± 2.9 |
| **ClpP.z4** | 0.64 ± 0.37 | 11.59 ± 7.20 | 0.12 ±0.03 | 3.04 ± 3.63 | 21.7 ± 2.9 |
| **DnaJ.tt** | 0.87 ± 0.30 | 4.42 ± 2.04 | 0.11 ± 0.02 | 0.56 ± 0.33 | 21.7 ± 2.9 |
| **DnaJ.z4** | 0.02 ± 0.04 | 0.95 ± 0.98 | 0.13 ± 0.02 | 2.35 ± 0.16 | 11.7 ± 2.9 |
| **DnaJ.jz** | 1.20 ± 0.10 | 6.97 ± 4.91 | 0.09 ±0.02 | 2.9 ± 0.71 | 11.7 ± 2.9 |
